# Generative modeling and latent space arithmetics predict single-cell perturbation response across cell types, studies and species

**DOI:** 10.1101/478503

**Authors:** M. Lotfollahi, F. Alexander Wolf, Fabian J. Theis

## Abstract

Accurately modeling cellular response to perturbations is a central goal of computational biology. While such modeling has been proposed based on statistical, mechanistic and machine learning models in specific settings, no generalization of predictions to phenomena absent from training data (‘out-of-sample’) has yet been demonstrated. Here, we present scGen, a model combining variational autoencoders and latent space vector arithmetics for high-dimensional single-cell gene expression data. In benchmarks across a broad range of examples, we show that scGen accurately models dose and infection response of cells across cell types, studies and species. In particular, we demonstrate that scGen learns cell type and species specific response implying that it captures features that distinguish responding from non-responding genes and cells. With the upcoming availability of large-scale atlases of organs in healthy state, we envision scGen to become a tool for experimental design through *in silico* screening of perturbation response in the context of disease and drug treatment.

## Introduction

Single-cell transcriptomics has become an established tool for unbiased profiling of complex and heterogeneous systems [1, 2]. The generated datasets are typically used for explaining phenotypes through cellular composition and dynamics. Of particular interest is the dynamics of single cells in response to perturbations, be it to dose [3], treatment [4, 5] or knock-out of genes [6–8]. Although advances in single-cell differential expression analysis [9, 10] enabled the identification of genes associated with a perturbation, generative modeling of perturbation response takes a step further in that it enables *in silico* generation of data. The ability of generating data that cover phenomena not seen during training, is particularly useful and referred to as ‘out-of-sample’ prediction.

While dynamic mechanistic models have been suggested for predicting low-dimensional quantities that characterize cellular response [11, 12], such as a scalar measure of proliferation, they face fundamental problems. These models cannot be easily formulated in a data-driven way and require temporal resolution of the experimental data. Due to the typically small number of time points available, parameters are often hard to identify. Resorting to linear statistical models for modeling perturbation response [6, 8], by contrast, leads to small predictive power for the complicated nonlinear effects that single-cell data display. By contrast, neural network models do not face these limits.

Recently, such models have been suggested for the analysis of single-cell RNA-seq data [13–17]. In particular, generative adversarial networks (GANs) have been proposed for simulating single cell differentiation through so a called latent space interpolation [16]. While being an interesting alternative to established pseudotemporal ordering algorithms [18], this analysis does not demonstrate the GAN’s capability of out-of-sample prediction. The use of GANs for the harder task of out-of-sample prediction is hindered by fundamental difficulties: (1) GANs are hard to train for structured high-dimensional data, leading to high-variance predictions with large errors in extrapolation, and (2), GANs do not allow to directly map a gene expression vector *x* on a latent space vector *z*, making it hard to impossible to generate a cell with wished properties. In addition, GANs for structured data have not yet shown advantages over the simpler variational autoencoders (VAE) [19] (Supplemental Note 1.1).

To overcome the problems inherent to GANs, we built scGen based on a VAE combined with vector arithmetics with an architecture adapted for single-cell RNA-seq data. For the first time, scGen enables predictions of dose and infection response of cells for phenomena absent from training data across cell types, studies and species. In a broad benchmark, it outperforms other potential modeling approaches such as linear methods, conditional variational autoencoders and style-transfer GANs. The benchmark of several generative neural network models should present a valuable resource for the community showing opportunities and limitations for such models when applied to transcriptomic data. scGen is based on Tensorflow [20] and on the single-cell analysis toolbox Scanpy [21].

## Results

### scGen accurately predicts single-cell perturbation response out-of-sample

High-dimensional scRNA-seq data is typically assumed to be well-parametrized by a low-dimensional manifold arising from the constraints of the underlying gene regulatory networks. Current algorithms mostly focus on characterizing the manifold using graph-based techniques [24, 25] in the space spanned by a few principal components. More recently, the manifold has been modeled using neural networks [13–17]. As in other application fields [26, 27], in the latent spaces of these models, the manifolds display astonishingly simple properties, such as approximately linear axes of variation for latent variables explaining a major part of the variability in the data. Hence, linear extrapolations of the low-dimensional manifold could in principle capture variability related to perturbation and other covariates (Supplemental Note 1.2, Supplemental Figure 1).

Let every cell *i* with expression profile *x*_*i*_ be characterized by a variable *p*_*i*_, which represents a discrete attribute across the whole manifold, such as perturbation, species or batch. To start with, we assume only two conditions 0 (unperturbed) and 1 (perturbed). Let us further consider the conditional distribution *P* (*x*_*i*_ *| z*_*i*_, *p*_*i*_), which assumes that each cell *x*_*i*_ comes from a low-dimensional representation *z*_*i*_ in condition *p*_*i*_. We use a VAE to model *P* (*x*_*i*_ *| z*_*i*_, *p*_*i*_) in its dependence on *z*_*i*_ and vector arithmetics in the VAE’s latent space to model the dependence on *p*_*i*_ (Figure 1).

**Figure 1.**
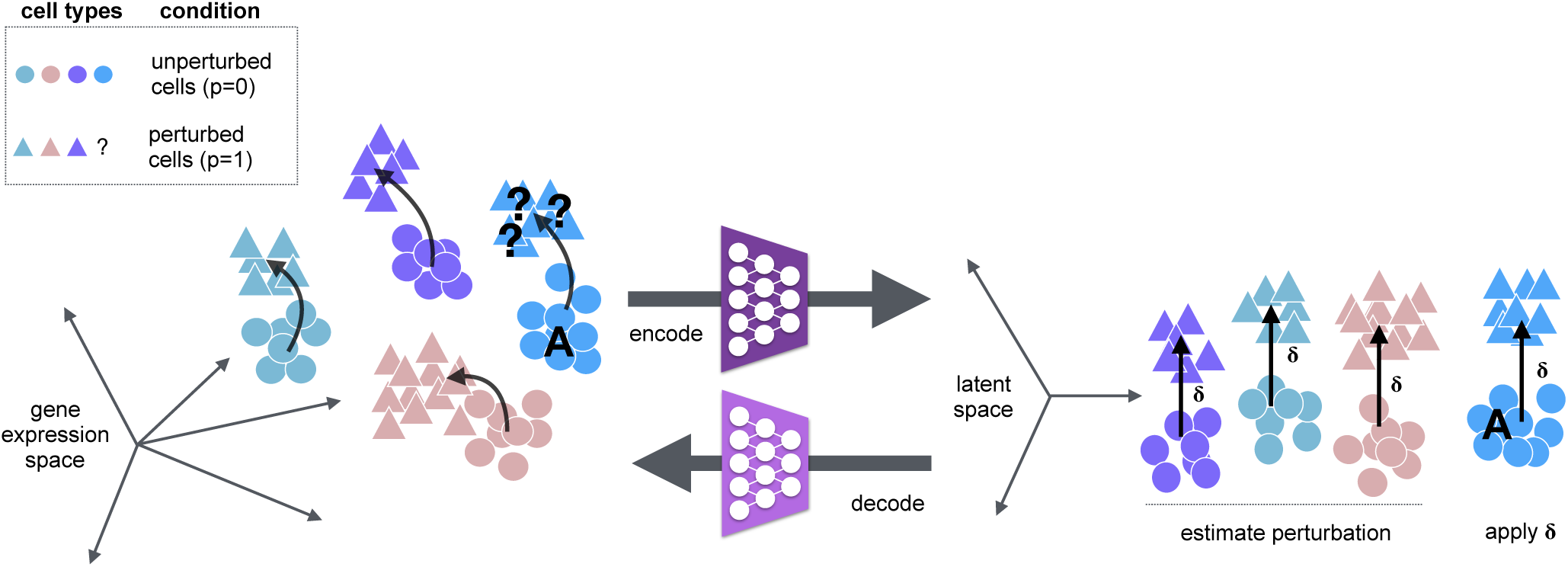
scGen, a method to predict single cell perturbation response. Given a set of observed cell types in control and stimulation, we aim to predict the perturbation response of a new cell type A (blue) by training a model that learns to generalize the response of the cells in the training set. Within scGen, the model is a variational autoencoder and the predictions are obtained using vector arithmetics in the autoencoder’s latent space. Specifically, we project gene expression measurements into a latent space using an encoder network and obtain a vector *δ* that represents the difference between perturbed and unperturbed cells from the training set in latent space. Using *δ*, unperturbed cells of type A are linearly extrapolated in latent space. The decoder network then maps the linear latent space predictions to highly non-linear predictions in gene expression space.

Equipped with this, consider a typical extrapolation problem. Assume cell type *A* exists in the training data only in the unperturbed (*p* = 0) condition. From that, we predict the latent representation of perturbed cells (*p* = 1) of cell type *A* using 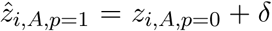, where *z*_*i,A,p*=0_ and 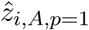 denotes the latent representation of cells with cell type *A* in conditions *p* = 0 and *p* = 1, respectively and *δ* is the difference vector of means between cells in the training set in condition 0 and 1 (Supplemental Note 1.3). From the latent space, scGen maps predicted cells to high-dimensional gene expression space using the generator network estimated while training the VAE.

To demonstrate the performance of scGen, we apply it to published human PBMC samples in control and under IFN-*β* stimulation [3] (Supplemental Note 2). As a first test, we compare the predictions of stimulated CD4-T cells held out during training (Figure 2a). scGen prediction of the mean associated with the perturbation in CD4-T cells correlates well with the ground-truth across all genes (Figure 2b). Comparing upregulated genes in stimulation (for example labeled transcripts in Figure 2c) we observe that these genes very well coincide in real and predicted stimulated cells. To evaluate generality, we trained six other models while holding out each of the six major cell types present in the study. Figure 2d shows that our model accurately predicts all other cell types (average *R*^2^ = 0.954). Moreover, the distribution of the strongest regulated IFN-*β* response gene *ISG15* as predicted by scGen not only provides a good estimate for the mean but also captures the variance of the distribution (Figure 2e, all genes in Supplemental Figure 2a).

**Figure 2.**
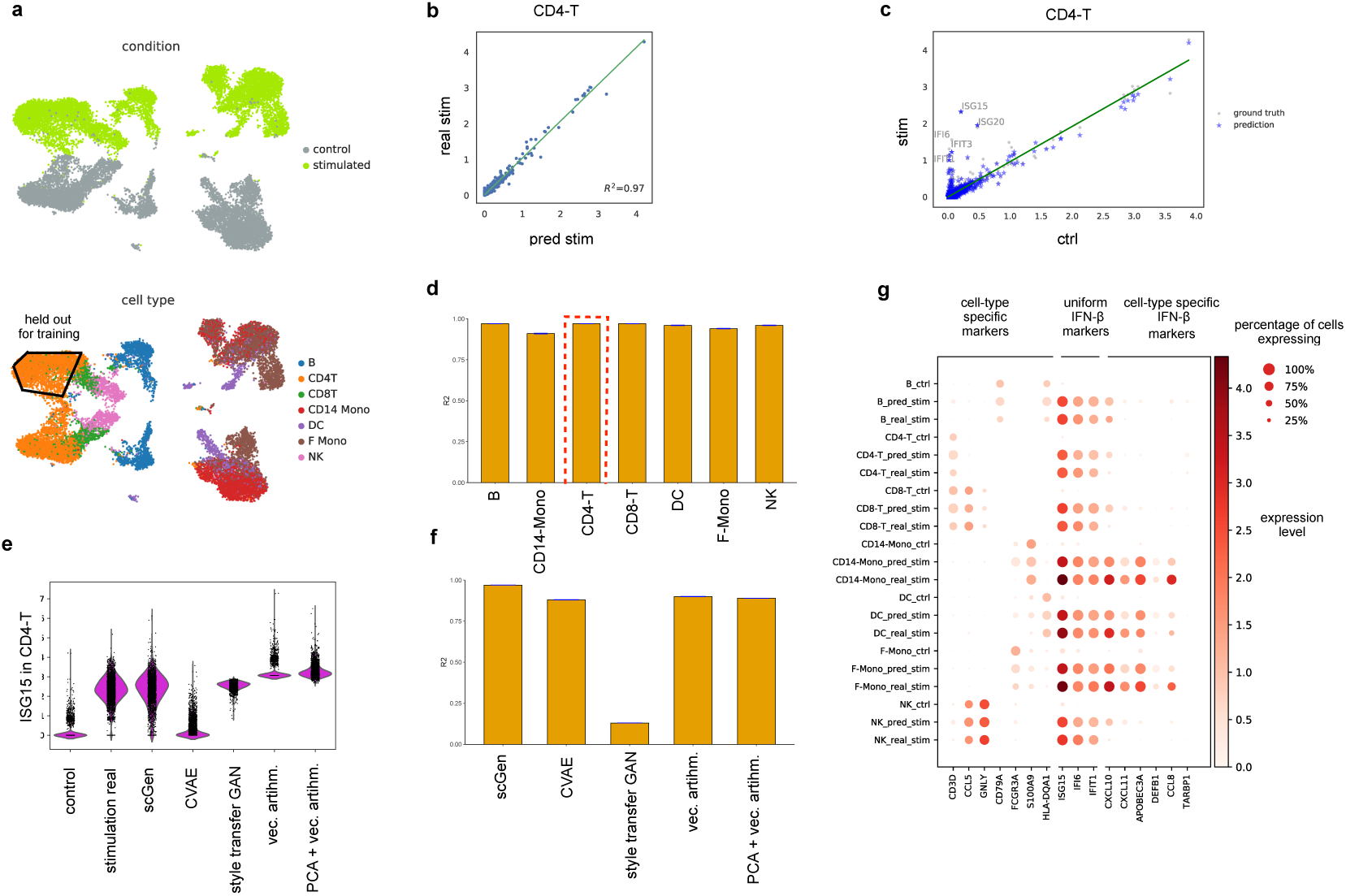
scGen accurately predicts single-cell perturbation response out-of-sample. **a,** UMAP visualization [22] of the distributions of condition, cell type and data split for the prediction of IFN-*β* stim-ulated CD4-T cells from altogether 16,893 PBMCs from Kang *et al.* [3]. **b,** Mean gene expression of 6,998 genes between scGen predicted and real stimulated CD4-T cells. **c,** Mean gene expression for control versus stimulated resp. predicted CD4-T cells together with top five upregulated differentially expressed genes. **d,** Comparison of *R*^2^ values for mean gene expression between real and predicted cells for the 7 different cell types of the study. **e,** Distribution of *ISG15*: the top uniform marker (response) gene to IFN-*β* [23] between control, predicted and real stimulated cells of scGen when compared with other potential prediction models. **f,** Similar comparison of *R*^2^ values to predict unseen CD4-T stimulated cells. **g,** Dot plot for comparing control, true and predicted stimulation when predicting on seven cell types from Kang *et al.*.

### scGen outperforms alternative modeling approaches

Aside from scGen, we studied further natural candidates for modeling a conditional distribution that is able to capture perturbation response. We benchmark scGen against four of these candidates, including two generative neural networks and two linear models. The first of these models is the conditional variational autoencoder (CVAE) (Supplemental Note 3, Supplemental Figure 3a, [28]), which has recently been adapted to preprocessing, batch-correcting and differential testing of single-cell data [13]. However, it has not been shown to be a viable approach for out-of-sample predictions, even though, formally, it readily admits the generation of samples from different conditions. The second class of models are style transfer GAN (Supplemental Note 4, Supplemental Figure 3b), which are commonly used for unsupervised image to image translation [29, 30]. In our implementation, such a model is directly trained for the task of transferring cells from one condition to another. The adversarial training is highly flexible and does not require an assumption of linearity in a latent space. In contrast to other propositions for mapping biological manifolds using GANs [31], style transfer GANs are able to handle unpaired data, a necessity for their applicability to single-cell RNA-seq data. We also mention that we tested ordinary GANs combined with vector arithmetics similar to Ghahramani *et al.* [16]. However, for the fundamental problems outlined above, we were not able to produce any meaningful out-of-sample predictions using this setup. In addition to the non-linear generative models, we tested simpler linear approaches based on vector arithmetics in gene expression space and the latent space of principal component analyses (PCA).

Applying the competing models to the PBMC dataset, we observe that all other models fail to predict mean and variance of the distribution of *ISG15* (all genes in Supplemental Figure 2), in stark contrast to scGen’s performance (Figure 2e). CVAE and style transfer GANs predictions are vaguely correlated with ground truth values and linear models also yield incorrect negative values (Supplemental Figure 2b-d). However, as shown in Figure 2b scGen provides most faithful prediction to real CD4-T cells and outperforms all other potential models (Figure 2f, Supplemental Figure 2, Supplemental Note 5).

A likely reason for why CVAE fails to provide meaningful out-of-sample predictions, is that it disentangles perturbation information from the latent space. Hence, the model does not learn non-trivial patterns linking perturbation to cell type. A likely reason for that the style transfer GAN is incapable for achieving the task is it’s attempt of matching two high-dimensional distributions, with much more complex models involved than in the case of scGen. While notoriously more difficult to train. Some of these arguments can be better understood when inspecting the latent space distribution embeddings of the generative models. As the CVAE completely strips off all perturbation-variation, its latent space embedding does not allow to distinguish perturbed from unperturbed cells (Supplemental Figure 4a). In contrast to CVAE representations, the scGen (VAE) latent space representation captures both information for condition and cell type (Supplemental Figure 4c), reflecting that non-trivial patterns across condition and cell type variability have been learned.

### scGen predicts both response shared among cell types and cell type specific response

Depending on shared or individual receptors, signaling pathways and regulatory networks, a group of cells perturbation response may result in expression-level changes that are shared across all cell types or unique to only some. Inferring both types of responses is essential for understanding mechanisms involved in disease progression as well as adequate drug dose predictions [32, 33]. Here, we show that scGen is able to capture both shared and cell type specific response after stimulation by IFN-*β* when any of the cell types in the data is held out during training and subsequently predicted (Figure 2g). For this, we use previously reported marker genes [23] of three different kinds: cell type specific markers independent of the perturbation such as *CD79A* for B cells, perturbation-response specific genes like *ISG15, IFI6, IFIT1* expressed in all cell types, and genes of cell type specific responses to the perturbation such as *APOBEC3A* in for DC cells. Across the seven different held out perturbed cell types present in the data of Kang *et al.*, scGen consistently makes good predictions not only of unperturbed and shared perturbation effects but also for cell type specific ones. Hence, although scGen encodes perturbation response by a shared *δ* across all cells in the latent space, after decoding to expression space both shared and individual changes can be captured.

### scGen robustly predicts intestinal epithelial cells response to infection

To illustrate that scGen works robustly, we evaluate its prediction performance quantitatively in two datasets from Haber *et al.* [4] related to epithelial cells from the small intestine (Supplemental Note 2) using the same network architecture as for the data of Kang *et al.*. These datasets consist of intestinal epithelial cells after *Salmonella* or *Heligmosomoides polygyrus (H.poly)* infections, respectively. scGen shows good performance for early transit-amplifying (TA.early) cells after infection with *H.poly* and *Salmonella* (Figure 3a,b), predicting both up and downregulated genes for each condition with high precision (*R*^2^ = 0.98 and *R*^2^ = 0.98, respectively). Figure 3c,d depicts similar analyses for both datasets and all occurring cell types — as before, the predicted ones being held out during training — indicating that scGen’s prediction accuracy is robust across most cell types. scGen’s performance is by far poorest for Tuft and Endocrine cells (Figure 3c,d). Whereas these cells, in reality, show a much weaker response than all other cells in the dataset, scGen predicts them as essentially non-responding (see Supplemental Figure 5). Hence, while scGen fails to capture the response quantitatively, it is remarkable that it captures the qualitative trend of the much weaker response despite not having seen this phenomenon for a high number of cells during training — both Endocrine and Tuft cells only constitute a small fraction of the data.

**Figure 3.**
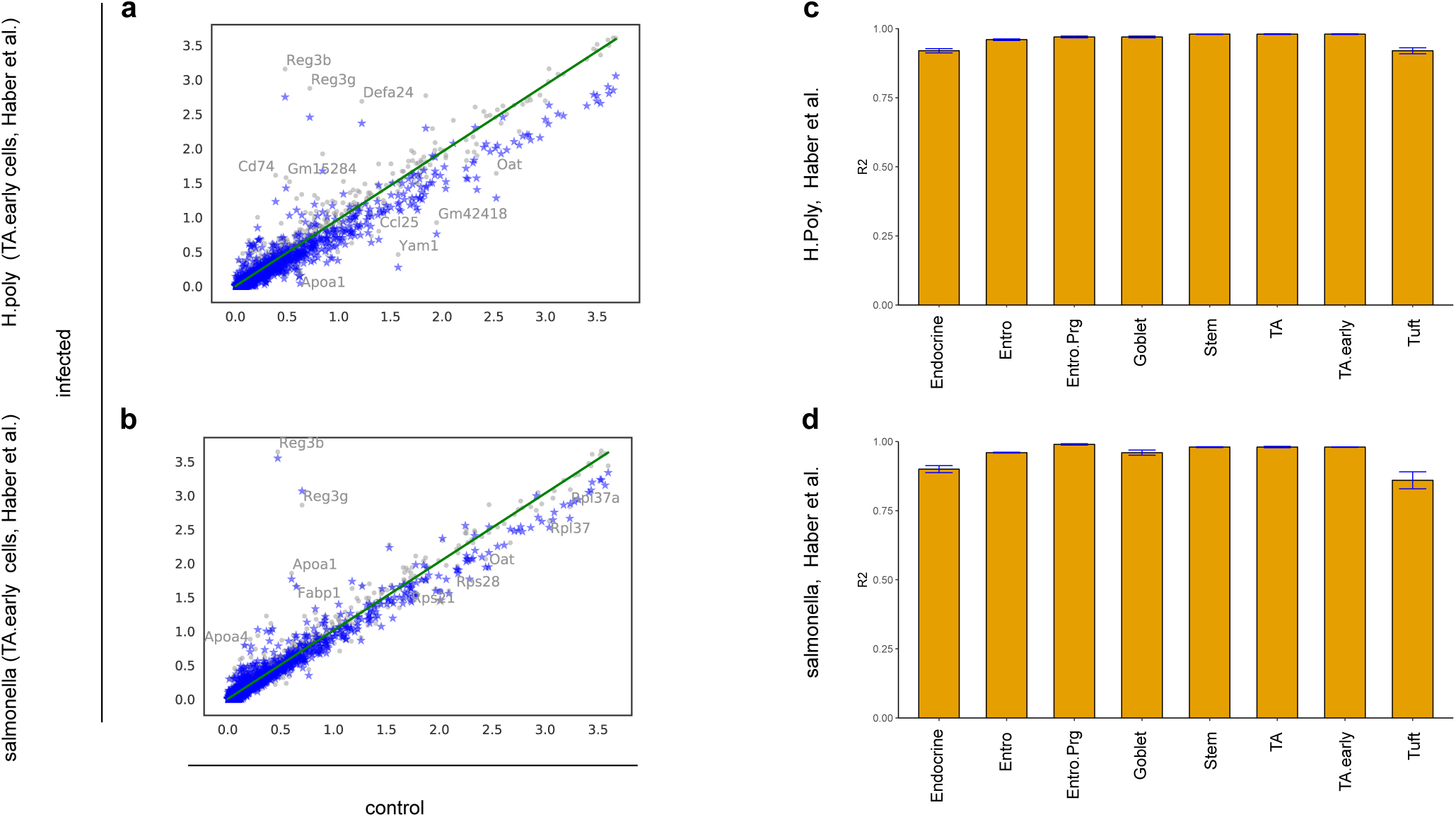
scGen models infection response in two datasets of intestinal epithelial cells. **a-b,** Prediction of early transit-amplifying (TA.early) cells from two different small intestine datasets from Haber *et al.* [4] infected with *Salmonella* and helminth *Heligmosomoides polygyrus (H.poly)* after 2 and 10 days, respectively. The mean gene expression for infected and control for different cell types shows how scGen transforms control to predicted perturbed cells in a way that the expression of top 5 up and downregulated differentially expressed genes are similar to real infected cells. **c-d,** Comparison of *R*^2^ values for mean gene expression between real and predicted cells for all the cell types in two different datasets illustrates that scGen performs well for all cell types in different scenarios.

In order to further understand when scGen starts to fail to make meaningful predictions, we again trained it on the PBMC data of Kang *et al.*, but now with more than one cell type held out. This study shows that scGen’s predictions are robust when holding out several dissimilar cell types (Supplemental Figure 6a-b) but start failing when training on data that only contains information about the response of one highly dissimilar cell type (see CD4-T predictions in Supplemental Figure 6c).

Finally, similar to what has been shown by [16] for differentiation of epidermal cells, we cannot only generate fully responding cell populations, but also intermediary cell states between two conditions. Here, we do so for the IFN-*β* stimulation and the *Salmonella* infection (Supplemental Note 6, Supplemental Figure 7).

### scGen enables cross-study predictions

We showed that scGen predicts cells from a cell type in a specific biological condition using all other cells available in that study. In order to be applicable to broad cell atlases such as the Human Cell Atlas [35], the algorithm ought to be robust against batch effects and hence generalize its prediction to unperturbed cells measured in a different study. For this, we consider a scenario with two single cell studies: study A, where cells within a specific organ have been observed in two biological conditions, e.g., control and stimulation, and study B with the same setting as study A but only in the control condition. By jointly encoding the two datasets, scGen provides a model for predicting the perturbation for study B (Figure 4a) by estimating the study effect as the linear perturbation in the latent space. To demonstrate this, we use as study A the PBMC dataset from Kang *et al.* and as study B another PBMC study consisting of 2623 cells that are available only in the control condition (Zheng *et al.* [34]). After training the model on data from study A, we use the trained model to predict how the PBMCs in study B would respond to stimulation with IFN-*β*.

**Figure 4.**
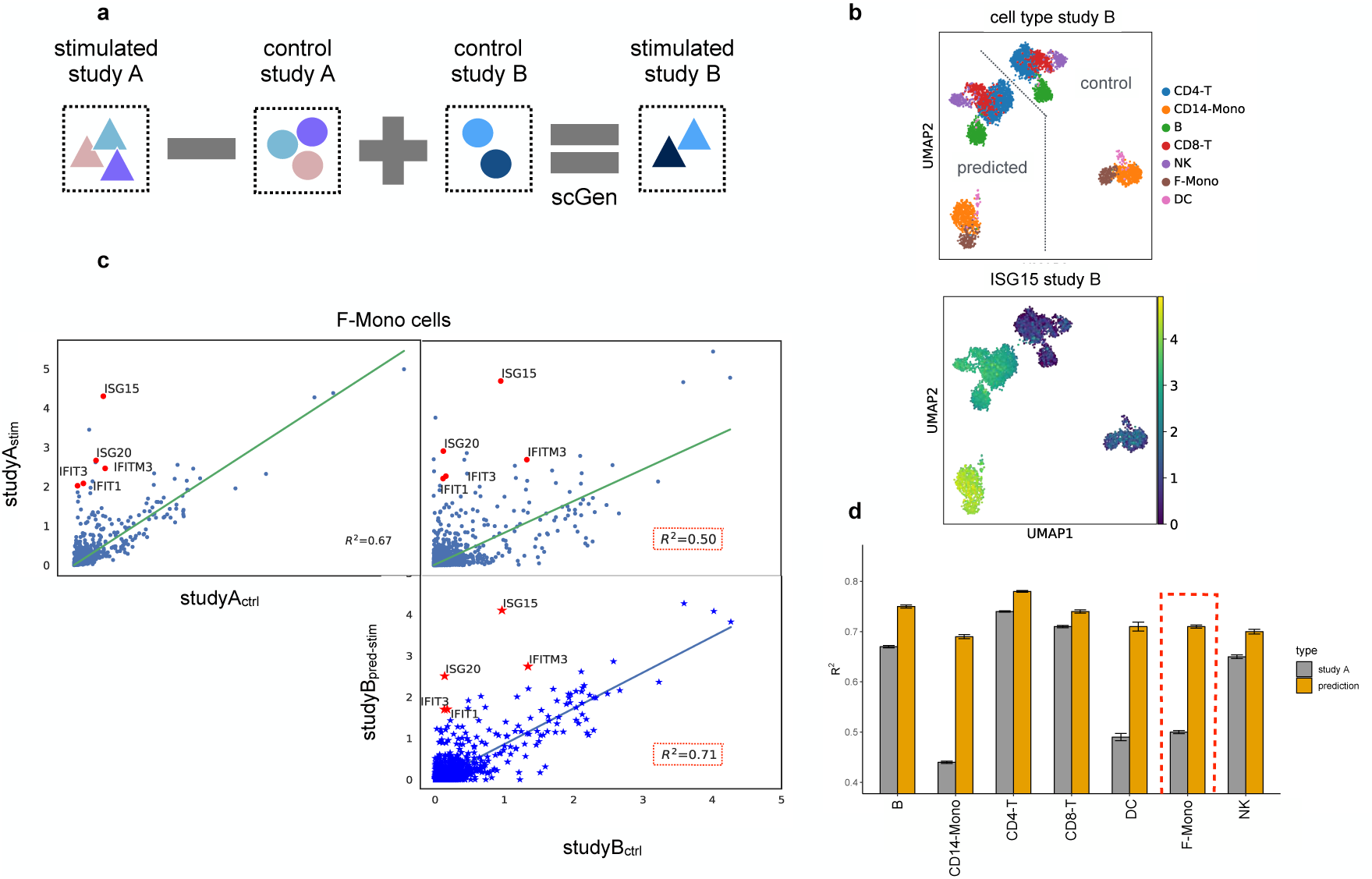
scGen accurately predicts single cell perturbation across different studies. **a,** scGen can be used to translate the effect of stimulation trained in study A to how stimulated cells in study B would look like, given a control sample set. **b,** Cell types for control and predicted stimulated cells for study B (Zheng *et al.* [34]) in two conditions where ISG15, the top IFN-*β* response gene, is only expressed in stimulated cells. **c,** Average expression between: control and stimulated F-Mono cells from study A (upper left), control from study B and stimulated cells from study A (upper right) and control from study B and predicted stimulated cells for study B (lower right). Red points denote top five differentially expressed genes for F-Mono cells after stimulation in study A. **d,** Comparison of *R*^2^ values highlighted in panel c for F-Mono and all other cell types.

As a first sanity check, we show that *ISG15* is also expressed in the prediction of stimulated cells based on the Zheng *et al.* (Figure 4b). This observation holds for all other differential genes associated with the stimulation, which we show for *FCGR3A+*-Monocytes (F-Mono) (Figure 4c): The predicted stimulated F-Mono cells correlate more strongly with the control cells in their study than with stimulated cells from study A while still expressing differentially expressed genes known from study A. Similarly, predictions for other cell types yield a higher correlation than the direct comparison with study A (Figure 4d).

### scGen predicts single-cell perturbation across species

In addition to learning the variation between two conditions, e.g. health and disease for a species, scGen can be used to predict across species. We trained a model on single cell RNA-seq dataset by Hagai *et al.* [5] comprised of bone marrow-derived mononuclear phagocytes from mouse, rat, rabbit, and pig perturbed with lipopolysaccharide (LPS) after six hours. Similar to what we did previously, we held out the rat LPS cells from the training data.

In contrast to previous scenarios, now, two global axes of variation exist in the latent space associated with species and stimulation, respectively.

Based on this, we have two latent difference vectors: *δ*_*LP* *S*_, which encodes the variation between control and LPS cells, and *δ*_*species*_, which accounts for differences between species. Next, we predict rat LPS cells using 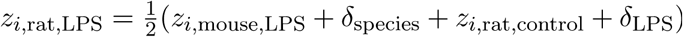. This equation takes an average of the two alternative ways of reaching rat LPS cells (Figure 5a). Figure 5(b) illustrates that predicted LPS cells express similar differential genes as true LPS stimulated rat cells. All other predictions along the major linear axes of variation also yield plausible results for stimulated rat cells (Supplemental Figure 8).

**Figure 5.**
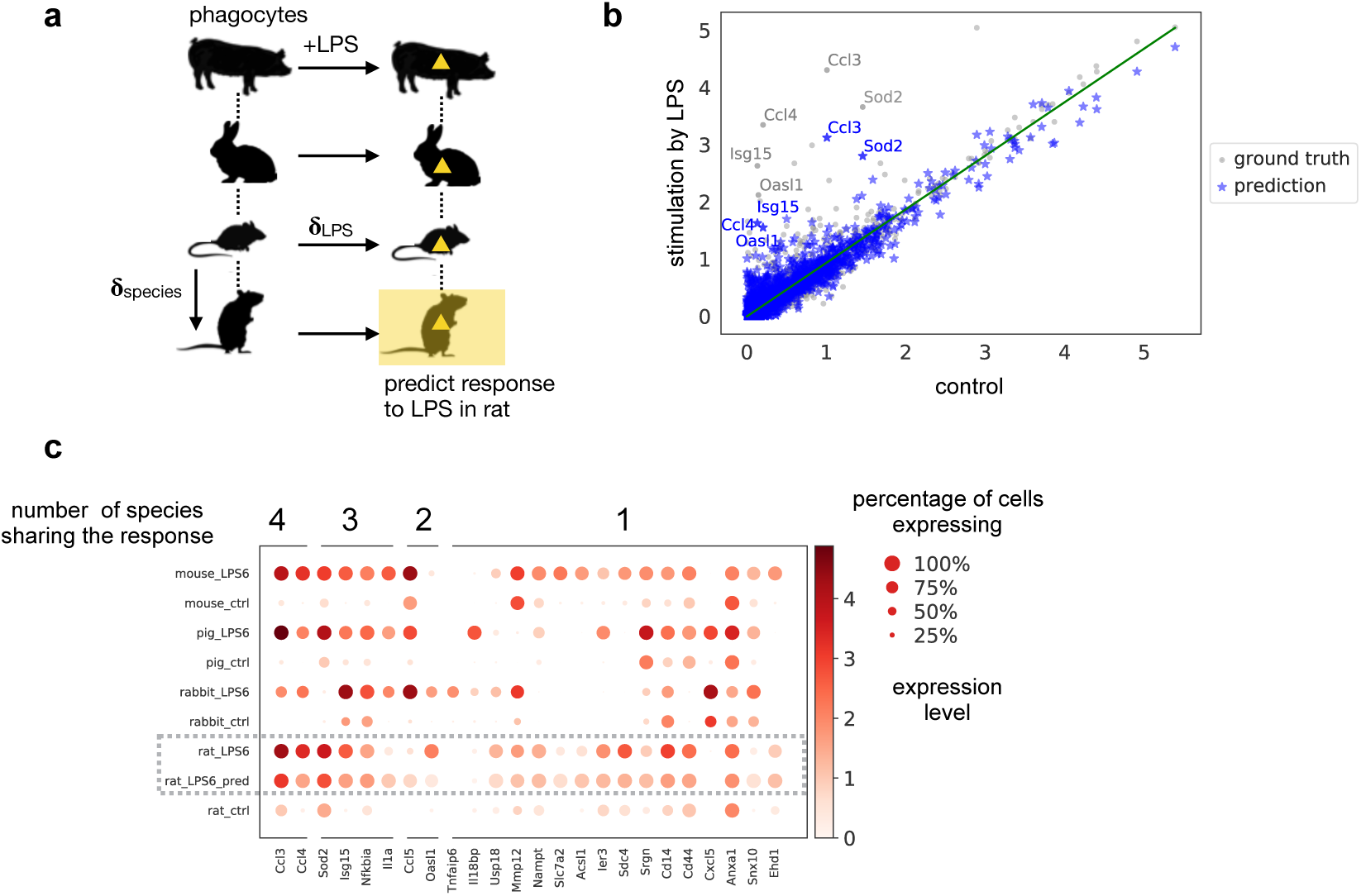
scGen predicts single cell perturbation response across different species. **a,** Prediction of unseen rat LPS phagocytes while accounting for both stimulation and species effect by learning two different vectors for each, on control and stimulated scRNA-seq from mouse, rabbit and pig by Hagai *et al.* [5]. **b,** Mean gene expression of 6,619 one-to-one orthologs between species for control rat cells plotted against true and predicted LPS while highlighted points represent top 5 differentially expressed genes after LPS stimulation in the real data. **c,** Dot plot of top 10 differentially expressed genes after LPS stimulation in each species, with numbers indicating how many species have those responsive genes among their top 10 differentially expressed genes.

In addition to the species-conserved response of a few upregulated genes, e.g. *Ccl3* and *Ccl4*, cells also display species specific responses. For example, *Il1a* is highly upregulated in all species except rat. Strikingly, scGen correctly identifies the rat cells as non-responding with this gene. Only the fraction of cells expressing *Il1a* increases at a low expression level (Figure 5c). Based on these early demonstrations, we foresee the prediction of human cell response based on data from healthy human and different healthy and perturbed animal models.

### scGen removes batch effects

Let us now show that scGen is able to efficiently correct for batch effects. To evaluate scGen’s batch correction capability, we merged four pancreatic datasets [36–39] (Figure 6a). We train scGen on these data and define a source and destination batch and compute a difference vector *δ*_*batch*_ between the source and the destination batch. To remove the batch effects from the destination batch, we add the learned *δ*_*batch*_ to the latent representation of the cells in the destination batch (Figure 6b). Using the cell type labels from the studies we observe a homogeneous overlap. A comparison with four existing batch removal methods (Supplemental Figure 9) shows that scGen performs as well as the other methods [23, 40–42]. To further evaluate batch removal ability of our model on a larger dataset, we merged eight different mouse single cell atlases comprised of 114600 cells from different organs [43–50]. As expected, the homogeneity of the data increased after batch correction (Supplemental Figure 10).

**Figure 6.**
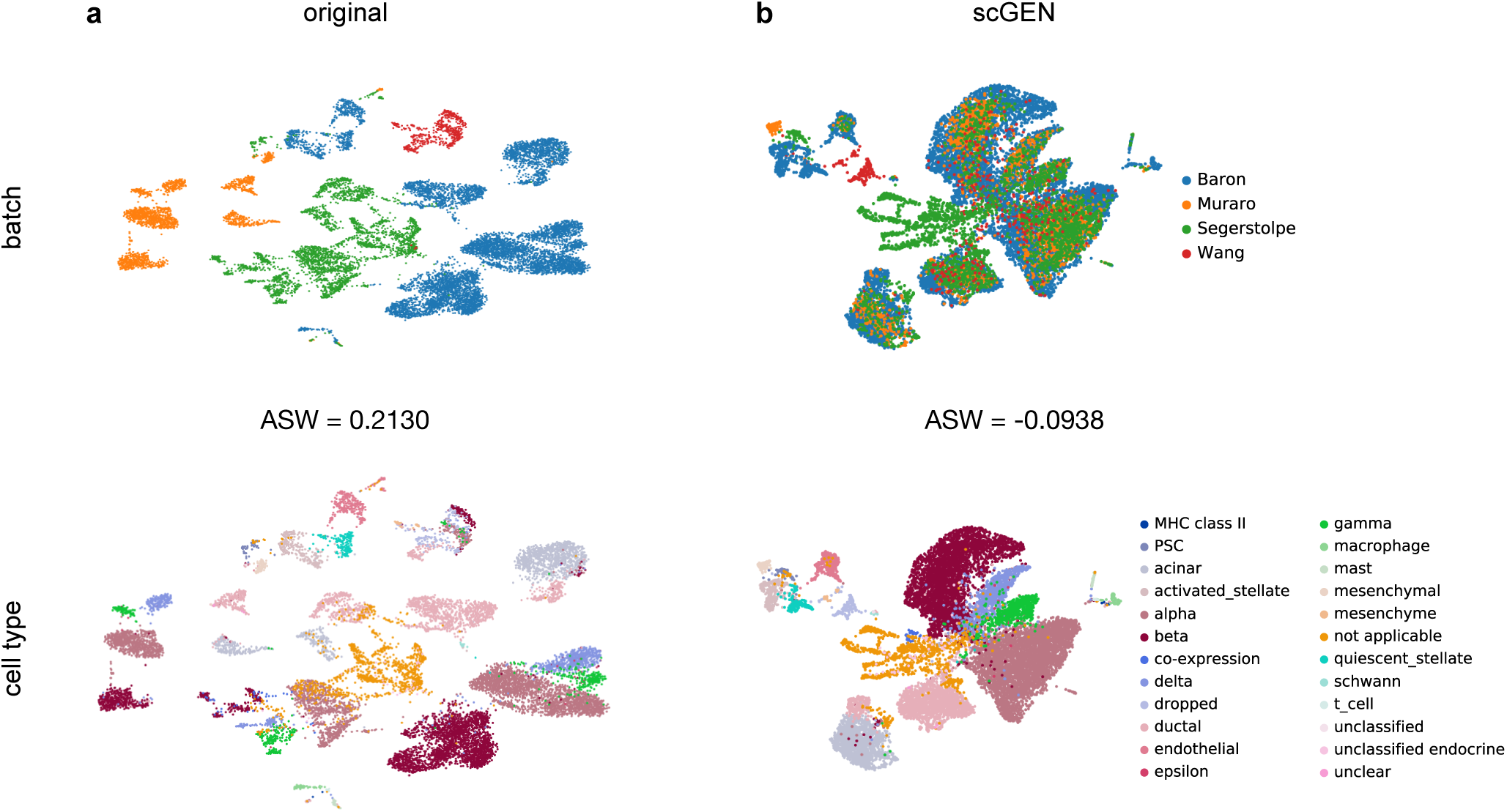
scGen removes batch effects. **a,** UMAP visualization of 4 technically diverse pancreatic datasets with their corresponding batch and cell types. We report average silhouette width (ASW) for batches in the original data (ASW = 0.2130, lower is better for batch effect evaluation). **b,** Data corrected by scGen mixes shared cell types from different studies while preserving study specific cell types independent (ASW = −0.0938).

## Discussion

We presented scGen, a model for predicting perturbation response of single cells based on generative neural networks and latent-space vector arithmetic. By adequately encoding the original expression space in a latent space, we achieve simple, near-to-linear mappings for highly non-linear sources of variation in the original data, which explain a large portion of the variability in the data. We provided examples for variation due to perturbation, species or batch. This allows to use scGen in several contexts including perturbation prediction response for unseen phenomena across cell types, study and species, for interpolating cells between conditions and for batch effect removal.

While we showed proof-of-concept for *in silico* predictions of cell type and species specific cellular response, in the present work, scGen has been trained on relatively small datasets, which only reflect subsets of biological and transcriptional variability. While we demonstrated scGen’s predictive power in these settings, a trained model cannot be expected to be predictive beyond the domain of the training data. To gain confidence in predictions, one needs to make realistic estimates for prediction errors by holding out parts of the data with known ground truth that are representative for the task. It is important to realize that such a procedure arises naturally when applying scGen in an alternating iteration of experiments, retraining based on new data and *in silico* prediction. By design, such strategies are expected to yield highly performing models for specific systems and perturbations of interest. It is evident that such strategies could readily exploit the upcoming availability of large-scale atlases of organs in healthy state, such as the Human Cell Atlas [35].

In summary, we demonstrated that scGen is able to learn cell type and species specific response. To be able to do so, the model needs to capture features that distinguish weakly from strongly responding genes and cells. Building biological interpretations of these features, for instance, along the lines of Ghahramani *et al.* [16] or Way and Greene [51], could help in understanding the differences between cells that respond to certain drugs and cells that do not respond, which is often crucial for understanding patient response to drugs [52].

## Code availability

Code is available from https://github.com/theislab/scGen.

## Data availability

All data is available from the original publications and linked on https://github.com/theislab/scGen.

## Author Contributions

M.L. performed the research, implemented the models and analyzed the data. F.A.W. conceived the project with contributions from M.L. and F.J.T.. F.A.W. and F.J.T. supervised the research. All authors wrote the manuscript.

## Acknowledgments

We are grateful to all members of the Theis lab, in particular, D.S. Fischer for early comments on predicting across species. M.L. is grateful for valuable feedback of L. Haghverdi regarding batch-effect removal. F.A.W. acknowledges discussions with N. Stranski on responding and non-responding cells and support by the Helmholtz Postdoc Programme, Initiative and Networking Fund of the Helmholtz Association. F.J.T. gratefully acknowledges support by the Helmholtz Association within the project “Sparse2Big” and by the German Research Foundation (DFG) within the Collaborative Research Centre 1243, Subproject A17.

During the work on the project, we became aware of reference [51], which suggests to study differences between cancer subtypes in the latent space of a VAE trained on bulk RNA-seq data from the Cancer Genome Atlas. The authors also demonstrate biological interpretability of these differences. In the weeks before submission of the manuscript, we became aware of the preprint [53], which addresses out-of-sample prediction in its revised version, but not in the context of single cell RNA-seq data.

## Supplemental Notes

### Supplemental Note 1: Models and theoretical background

#### Supplemental Note 1.1: Variational autoencoders

A variational autoencoder is a neural network consisting of an encoder and a decoder similar to classical autoencoders. Unlike the classical autoencoders, VAEs are able to generate new data points. The mathematics behind VAEs is not similar to classical autoencoders. The difference is that the model maximizes the likelihood of each sample *x*_*i*_ in the training set under a generative process as formulated in Equation (1).

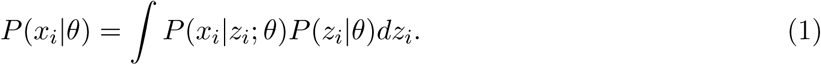

where *θ* is the model parameter which in our model corresponds to a neural network with its learnable parameters and *z*_*i*_ is a latent variable. The most important idea of a VAE is to sample latent variables *z*_*i*_ that are likely to produce *x*_*i*_ and using those to compute *P* (*x*_*i*_ *| θ*) [54]. We approximate the posterior distribution *P* (*z*_*i*_ *| x*_*i*_, *θ*) using the variational distribution *Q*(*z*_*i*_ *| x*_*i*_, *ϕ*) which is modeled by a neural network with parameter *ϕ*, called the inference network (the encoder). Next, we need a distance measure between the true posterior *P* (*z*_*i*_*|x*_*i*_, *θ*) and the variational distribution. To compute such a distance we use the Kullback-Leibler (𝕂 𝕃) divergence between *Q*(*z*_*i*_ *| x*_*i*_, *ϕ*) and *P* (*z*_*i*_ *| x*_*i*_, *θ*), which yields:

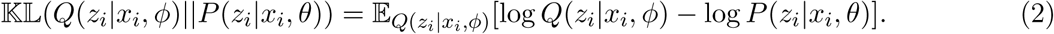

Now, we can derive both *P* (*x*_*i*_ *| θ*) and *P* (*x*_*i*_ *| z*_*i*_, *θ*) by applying Bayes rule to *P* (*z*_*i*_ *| x*_*i*_, *θ*) which results in:

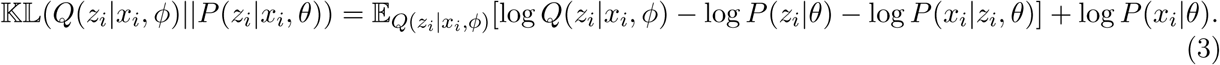

Finally, by rearranging some terms and exploiting the definition of KL divergence we have:

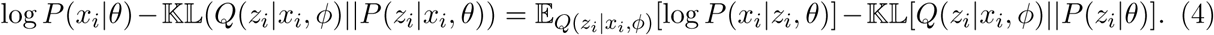

On the left hand side of Equation (4), we have the log-likelihood of the data denoted by log *P* (*x*_*i*_ *| θ*) and an error term which depends on the capacity of the model. This term ensures that *Q* is as complex as *P* and assuming a high capacity model for *Q*(*z*_*i*_*|x*_*i*_, *ϕ*), this term will be zero [54]. Therefore, we will directly optimize log *P* (*x*_*i*_*|θ*):

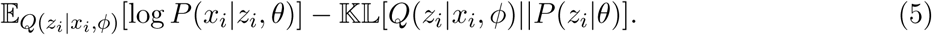

In order to maximize the Equation (5), we choose the variational distribution *Q*(*z*_*i*_*|x*_*i*_, *ϕ*) to be a multivariate Gaussian *Q*(*z*_*i*_*|x*_*i*_) = *𝒩* (*z*_*i*_; *µ*_*ϕ*_(*x*_*i*_), *Σ*_*ϕ*_(*x*_*i*_)) where *µ*_*ϕ*_ and *Σ*_*ϕ*_ are implemented with the encoder neural network and *Σ*_*ϕ*_ is constrained to be a diagonal matrix. The 𝕂 𝕃 term in Equation can be computed analytically since both both prior (*P* (*z*_*i*_ *| θ*)) and posterior (*Q*(*z*_*i*_ *| x*_*i*_, *ϕ*)) are multivariate Gaussian distributions. The integration for the first term in (5) has no closed-form and we need Monte Carlo integration to estimate it. We can sample *Q*(*z*_*i*_ *| x*_*i*_, *ϕ*) *L* times and directly use stochastic gradient descent to optimize Equation (6) as loss function for every training point *x*_*i*_ from dataset *D*:

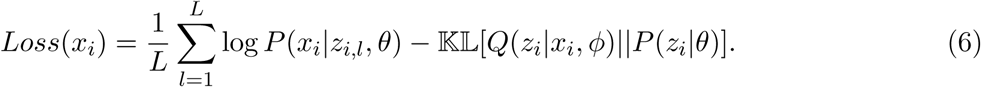

However, the first term in Equation (6) only depends on the the parameters of *P* and the parameters of variational distribution *Q* are not there. Therefore, it has no gradient with respect to *ϕ* to be back-propagated. In order to address this, the *reparameterization trick* [19] has been proposed. This trick works by first sampling from *∈ ∼𝒩* (0, *I*) and then computing 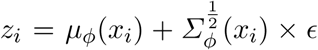. In consequence, we can use gradient-based algorithms to optimize Equation (6).

For the results shown in the present paper, we adapted the cost function (6) of the VAE by replacing *µ*(*x*_*i*_)^2^ with (log *Σ*(*x*_*i*_))^2^ in the regularization (𝕂𝕃) term.

#### Supplemental Note 1.2: Linearity of the latent space

scGen exploits vector arithmetics in the latent space of VAEs which assumes the shift (response) induced by stimuli can be modeled linearly. Similar to what has been shown by [55], we empirically demonstrate the linearity of the latent space with respect to biological conditions. In pursuance of that, we design a simple linear classifier based on the difference vector (*δ*) between two conditions in the latent space. We hypothesize that the *δ* vector directs toward a direction in the latent space where condition 1 increases. Therefore, by moving along the direction of *δ* we are moving from the condition 0 to condition 1. A high-level intuition for this is the difference vector manipulates cells by adding and removing information to them. Suppose, for example, a dimension of the latent vector corresponds to the degree of the infection in a cell. Increasing that attribute would be as easy as adding the *δ* vector corresponding for that attribute. In consequence, the dot product of the cells from the condition 1 with *δ* will be approximately greater than zero (or a constant positive value) indicating high similarity. Similarly, the dot product with cells in condition 0 would yield negative values showing low similarity (Supplemental Figure 1a). After finding the difference vector for each condition, including IFN-*β* from Kang *et al.* [3], *H.poly* and *Salmonella* infections from Haber *et al.* [4], we demonstrate the histogram of dot product results for the latent representation of all cells with their corresponding difference vector (Supplemental Figure 1b).

We did another test by calculating *δ*_stim-k_ denoting the difference between stimulated and control cells for cell type *k*. We also calculated another set of difference vectors, *δ*_celltype-ij_, representing the difference between each of the seven cell types present in Kang *et al.* dataset irrespective of their condition. Next, we calculated the cosine similarity for each set of previous vectors with *δ*. Supplemental Figure 1c depicts that vector in *δ*_stim-k_ set have very high cosine similarity with *δ* showing that they are both directing toward the same direction with a small angle. However, most of the *δ*_celltype-ij_ vectors have cosine similarity close to zero that shows the cell type and condition vectors are different and nearly orthogonal. In order to get an intuition of how unlikely is to get a high cosine similarity in 100-dimensional vector space, we randomly drew 1000 samples from 100- dimensional standard normal distribution and calculated pair-wise cosine similarity between them (Supplemental Figure 1c, random).

**Supplemental Figure 1.**
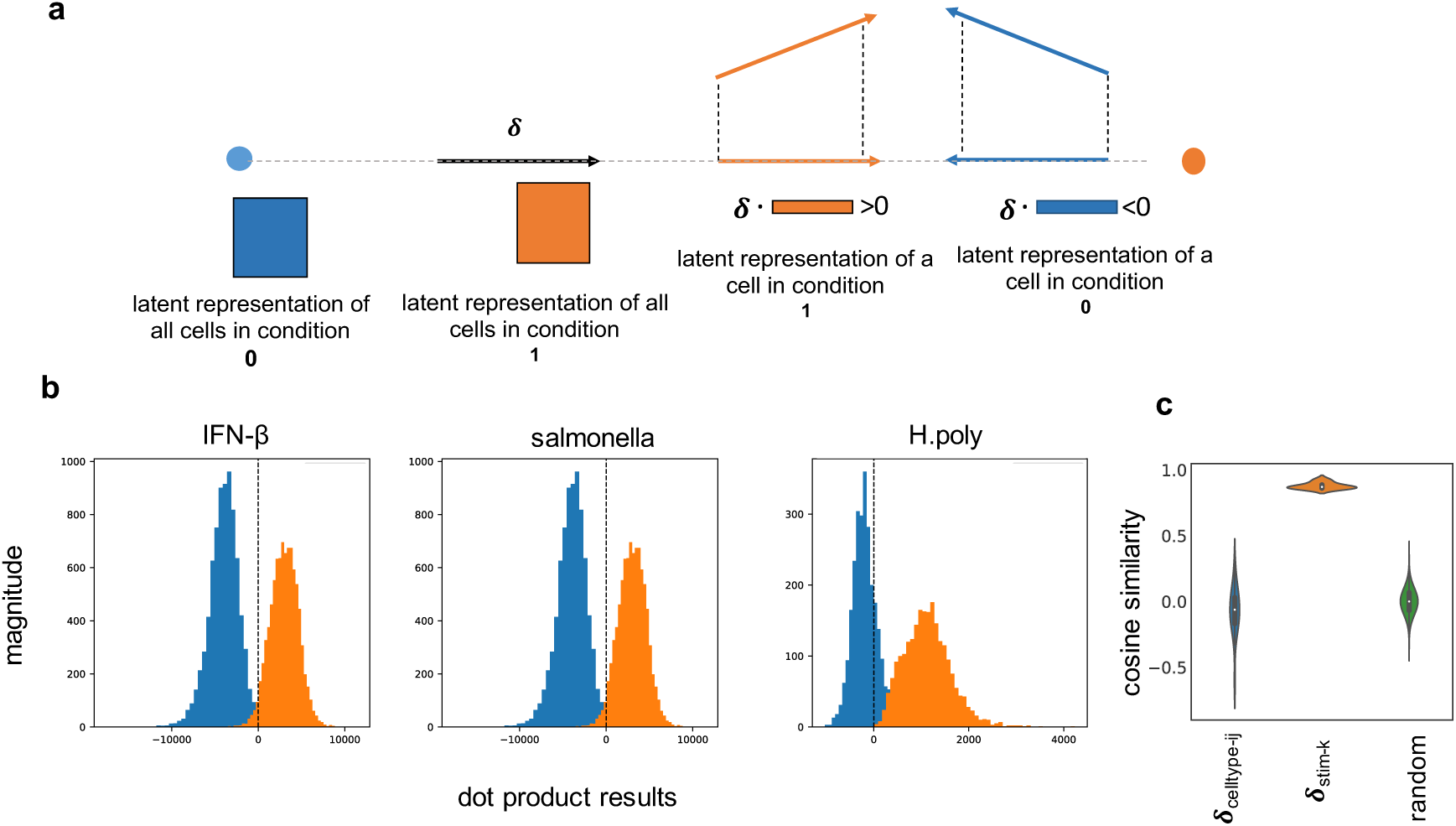
Linearity of the latent space. **a,** Building a linear classifier based on the dot product between the difference vector (*δ*) and the latent representation of each cell. **b,** Dot product results between latent representation of all cells with their corresponding difference vector (*δ*) for each condition shows that two conditions are approximately linearly separable using dot product classifier. **c,** Cosine similarity of *δ*_stim-k_, *δ*_celltype-ij_ with *δ* where *δ*_celltype-ij_ = avg(*z*_celltype_=*i*) − avg(*z*_celltype_=*j*) and *δ*_stim-k_ = avg(*z*_stim, celltype_=_k_) − avg(*z*_ctrl, celltype=k_) for all seven cell types present in Kang *et al.* dataset (*z* denotes the latent representation of all cells with the corresponding label). The third violin plot shows pairwise cosine similarity for a set of 1000 random samples from 100 dimensional standard normal distribution.

#### Supplemental Note 1.3: *δ* vector estimation

In order to estimate *δ*, first, we extract all cells for each condition. Next, for each cell type, we up-sample the cell type sizes to be equal to the maximum cell type size in that condition. To further remove the population size bias, we randomly down-sample the condition with a higher sample size to match the sample size of the other the condition. Finally, we estimate the difference vector by calculating *δ* = avg(z_condition=1_) *-* avg(z_condition=0_), where z_condition=1_ and z_condition=0_ denote the latent representation of cells in each condition, respectively.

#### Supplemental Note 2: Datasets

The First dataset includes two groups of peripheral blood mononuclear cells (PBMCs) from Kang *et al.* [3]. The original dataset includes 29065 cells split into 14446 stimulated and 14619 control cells from 8 individuals. We annotated cell types by extracting an average of top 20 cluster genes from each of 8 identified cell types in PMBCs from [34]. Next, the Spearman correlation between every single cell and all 8 cluster averages was calculated and each cell was assigned to the cell type which it had a maximum correlation (similar to [3]). After identifying cell types, Megakaryocyte cells were removed from the dataset due to the high uncertainty of assigned labels. Next, the dataset was filtered for cells with minimum 500 expressed genes and genes which were expressed at least in 5 cells. Moreover, we normalized counts per-cell and top 6998 differentially expressed genes were selected. Finally, we log-transformed the data in order to have a smoother training procedure.

**Supplemental Figure 2.**
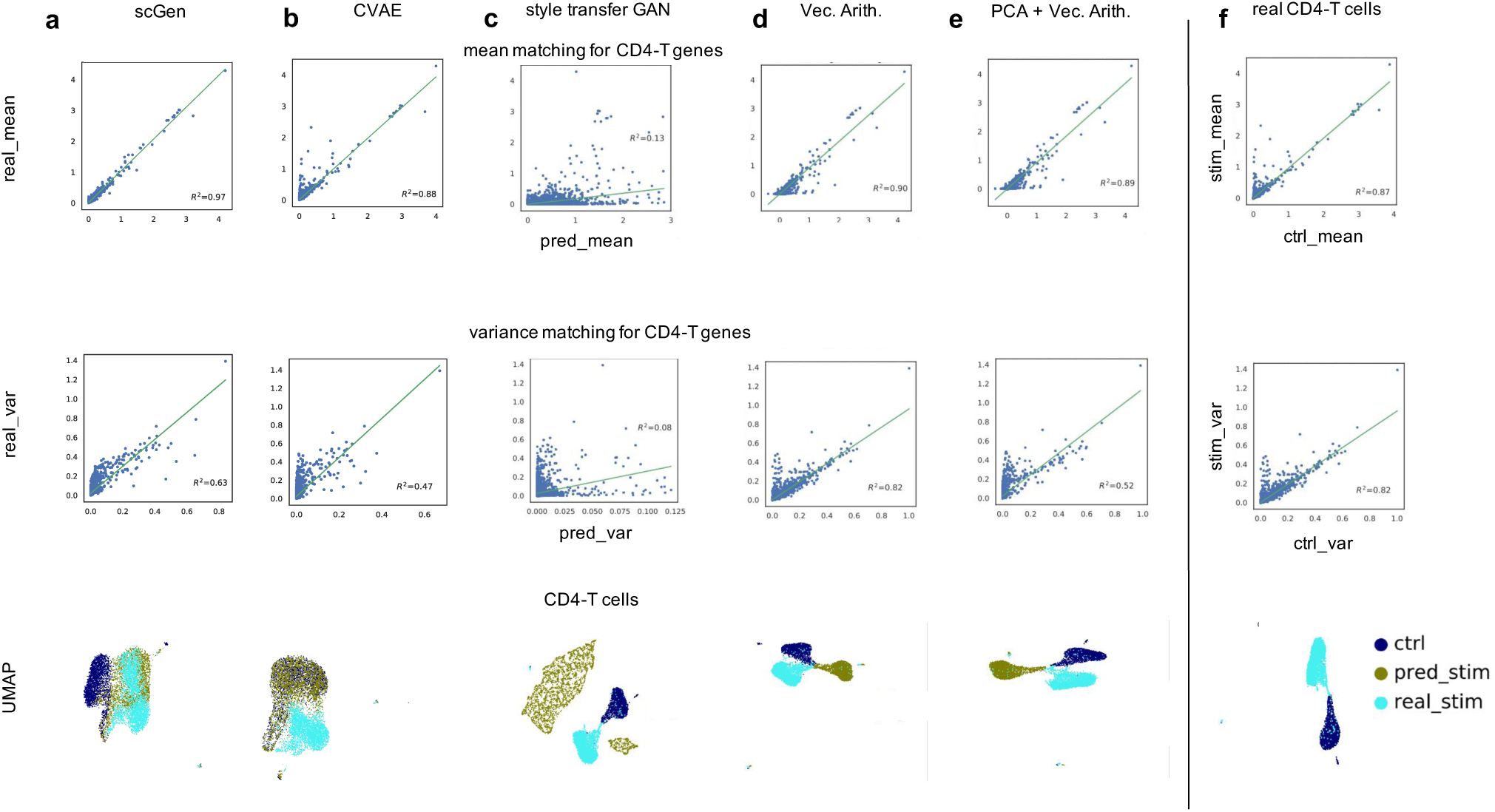
Distribution matching comparison between different models. **a-e,** Mean and variance matching comparison between scGen and four alternative models for CD4-T cells, shows scGen outperforms other models. Similarly, by comparing UMAP visualizations one can see predictions by scGen have more overlap with ground truth cells whereas predictions from other models lie far from real stimulated cells. **f,** Ground truth mean and variance between control and stimulated CD4-T cells.

The second dataset comprises of epithelial response to pathogen infection from Haber *et al.* [4]. In this dataset, the response of intestinal epithelial cells to *Salmonella* and parasitic helminth *Heligmosomoides polygyrus (H.poly)* were investigated. Moreover, it includes three different conditions including, 1777 *Salmonella* infected cells and ten days (2,711) after *H.poly* infection and finally a group of 3240 control cells. The data was normalized per-cell and top 7000 differentially expressed genes were selected and finally log-transformed.

The second PBMC dataset from Zheng *et al.* [34] was obtained from http://cf.10xgenomics.com/samples/cell-exp/1.1.0/pbmc3k/pbmc3k_filtered_gene_bc_matrices.tar.gz. After filtering cells, the data was merged with filtered PBMCs from Kang *et al.* [3]. The Megakaryocyte cells were removed from the smaller dataset. Next, the data was normalized and then we selected top 7000 differentially expressed genes. The merged dataset was log-transformed and cells from Kang *et al.* were used for training the model. The remaining 2623 cell from Zheng *et al.* were used for the prediction.

**Supplemental Figure 3.**
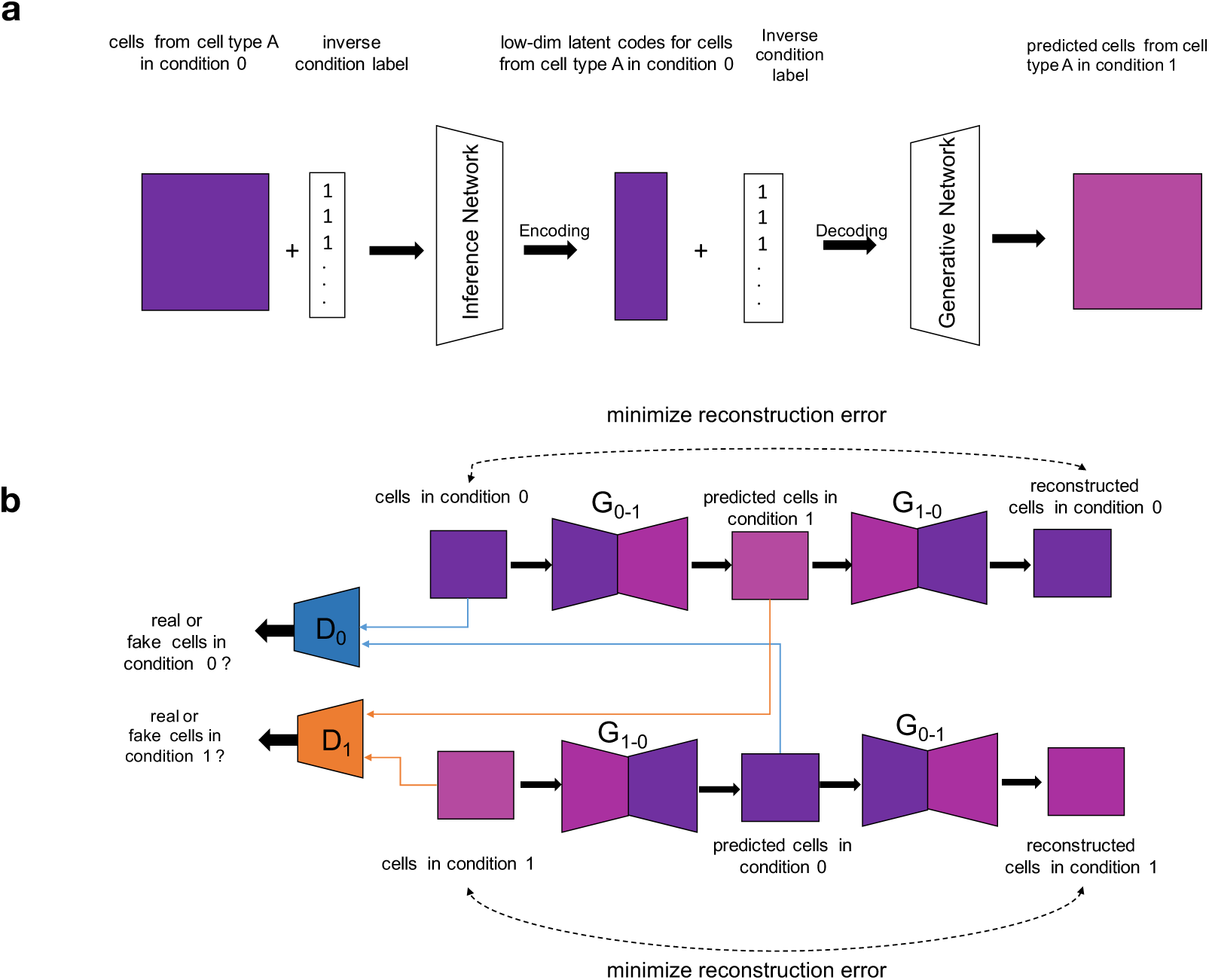
Graphical pipeline of two alternative approaches to predict unseen single cell perturbations. **a,** CVAE pipeline at test time to predict unseen condition. In order to predict cells in condition 1, we feed all cells present in condition 0 with inverse label 1 concatenated (shown with + symbol) to the data matrix. This informs the model that these cells are from condition 1. Therefore, the model changes the condition of input cells from 0 to 1. **b,** The style transfer GAN to transform one condition to another. This would be possible by learning a joint two-way mapping in an adversarial learning setting. There exist two generators, *G*_0−1_ which transforms cells from condition 0 to 1 and *G*_1−0_ which does the same task but in the reverse direction. Two discriminators, denoted by *D*_0_ and *D*_1_, are trained to detect real from fake cells generated by *G*_1−0_ and *G*_0−1_, respectively.

Pancreatic datasets were downloaded from ftp://ngs.sanger.ac.uk/production/teichmann/BBKNN/objects-pancreas.zip. All the comparisons to other batch corrections methods were performed similar to [41] with n = 50 PCs. The data was already preprocessed and directly used for training the model.

Mouse cell atlases were obtained from ftp://ngs.sanger.ac.uk/production/teichmann/BBKNN/ MouseAtlas.zip. The data was already preprocessed and directly used for training the model.

LPS dataset [5] was obtained from https://www.ebi.ac.uk/arrayexpress/experiments/E-MTAB-6754/?query=tzachi+hagai. The data were further filtered for cells, normalized and log-transformed. We used BiomaRt (v84) [56] to find ENSEMBL IDs of the 1-to-1 orthologs in the other three species with the mouse. In total 6619 genes were selected from all species for training the model.

**Supplemental Figure 4.**
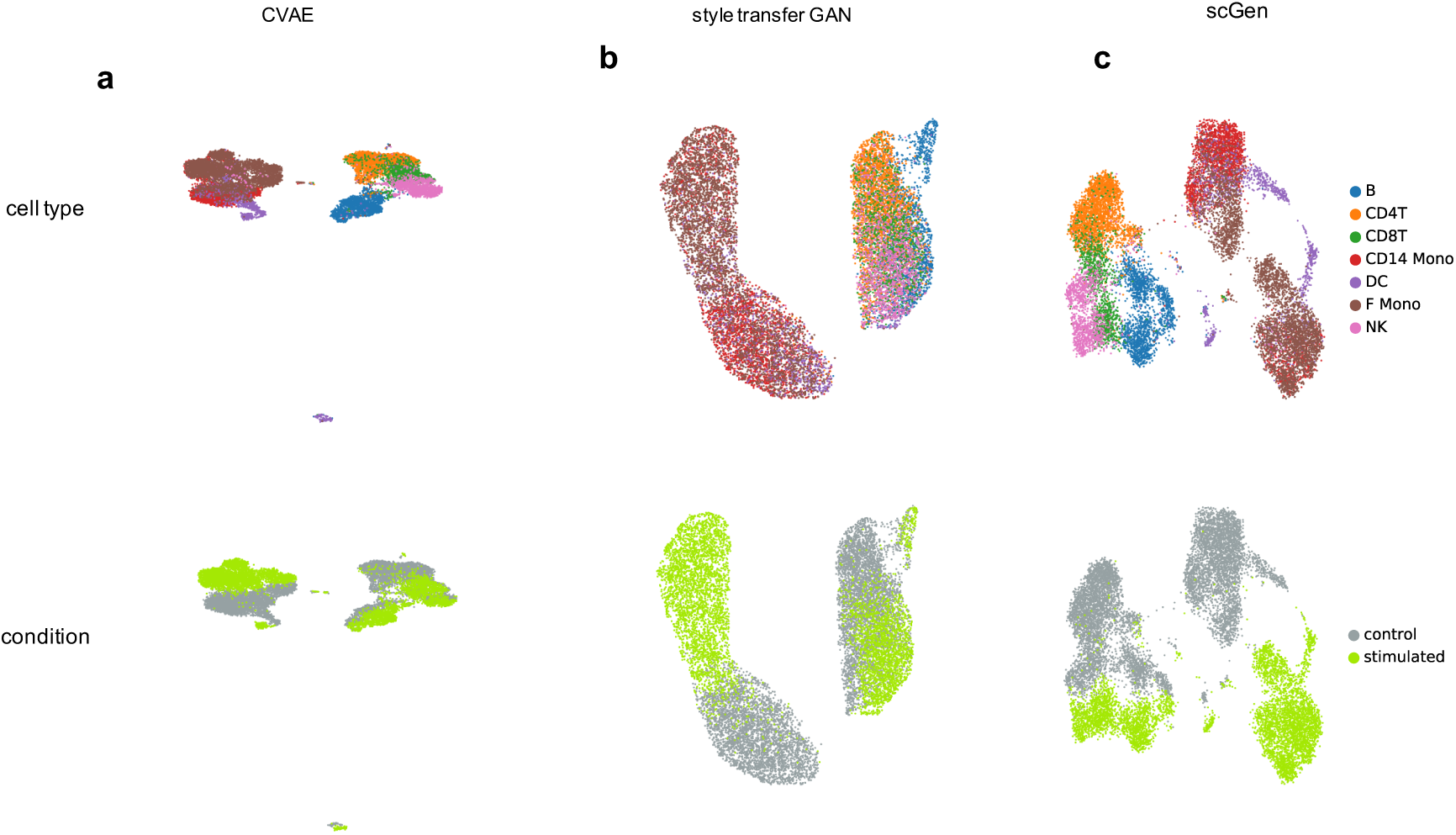
Latent space comparison. **a-c,** UMAP visualization of latent space representation for PBMCs from Kang *et al.* dataset. For scGen (VAE) and CVAE we used the bottleneck layer but for style transfer GAN we used discriminator’s penultimate output as the input for UMAP algorithm.

#### Supplemental Note 3: Conditional variational autoencoder

The conditional variational autoencoder (CVAE) [28] is also based on the variational inference framework. In the CVAE setting one can train a model conditioned on two existing biological conditions. We concatenate the condition of every cell with its input (*x*_*i*_) and latent variable (*z*_*i*_). At test time, we feed the model with cells in condition 0 and the label of condition 1 (inverse label) to transform the cells to same cell type but in condition 1 (Supplemental Figure 3**a**).

#### Supplemental Note 4: Style transfer GAN

The original style transfer model [30] learns to transform images in one visual domain (e.g., domain of all horses) to another domain (e.g., the domain of all zebras). We can adapt this to the single cell domain by training a network that receives single cells in condition 0 and transforms them to similar cells with the same cell type but in condition 1. This can be achieved in an adversarial training fashion (Supplemental Figure 3**b**). As it is shown in Supplemental Figure 3**b**, the model transforms cells in condition 0 to cells in condition 1 via *G*_0*–*1_ and then transforms them back to condition 1 using *G*_1*–*0_. There exists a second line of networks which learns to transform cells from condition 1 to 0 and reconstruct them back to condition 0. These two pipelines must work in a way that they can fool two discriminators (one for each condition) which are trained to detect real cells from generated (fake) cells. In order to make the problem setting more constrained, the reconstructions should not highly deviate from the real data according to a distance metric (e.g., *L*_2_). Moreover, similar networks in both lines share parameters. At test time, one can feed the gene expression profile of all target cells in condition 0 to transform them to condition 1.

**Supplemental Figure 5.**
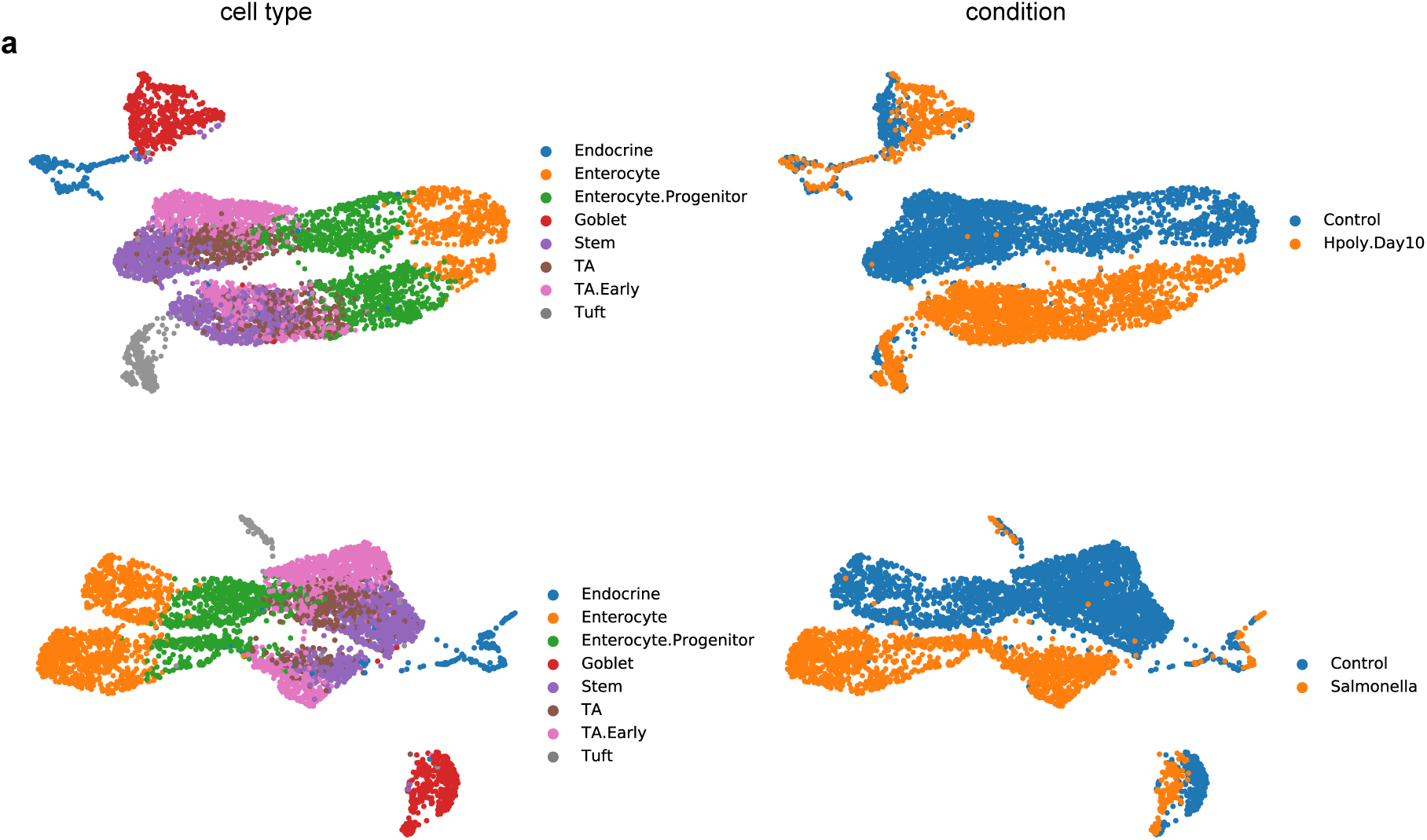
UMAP visualization for epithelial response to pathogen infection from Haber *et al.* [4]. **a,** Different cell types have various degree of response after infection. In comparison with other cell types, the Endocrine and Tuft cells are less affected after infection.

#### Supplemental Note 5: Model comparison

We compare the distribution matching capability of each model based on their variance and mean estimation of every individual gene. Our model yields most accurate mean estimation (*R*^2^ = 0.97, Supplemental Figure 2a) while other models yield poor results. For example, CVAE completely fails to upregulate differentially expressed genes and the result is more similar to control cells (*R*^2^ = 0.88, Supplemental Figure 2b). Notably, applying vector arithmetics in gene expression and PCA space make the mean of some genes to take invalid negative values and leaves the variance intact as it was in the real control cells (Supplemental Figure 2d,e). Furthermore, scGen also show reasonable performance in variance estimation (*R*^2^ = 0.63) and outperforms all other models (Supplemental Figure 2a).

#### Supplemental Note 6: Latent space interpolation

We exemplify the latent space interpolation ability of our model by generating 2000 intermediary TA (*Salmonella*, Haber *et al.*) and CD4-T (IFN-*β*, Kang *et al.*) cells. First, we project average control and predicted cells into the latent space and then linearly interpolate 2000 intermediary points between them. Next, by using generator network we map back latent intermediary cells into high-dimensional gene expression space (Supplemental Figure 7a-b). One can observe a smooth change of the top five up and downregulated *Salmonella* response genes as we traverse cell manifold from control towards *Salmonella* cells (Supplemental Figure7c). Similarly, we can see the upregulation of top five IFN-*β* response genes (Supplemental Figure 7d).

**Supplemental Figure 6.**
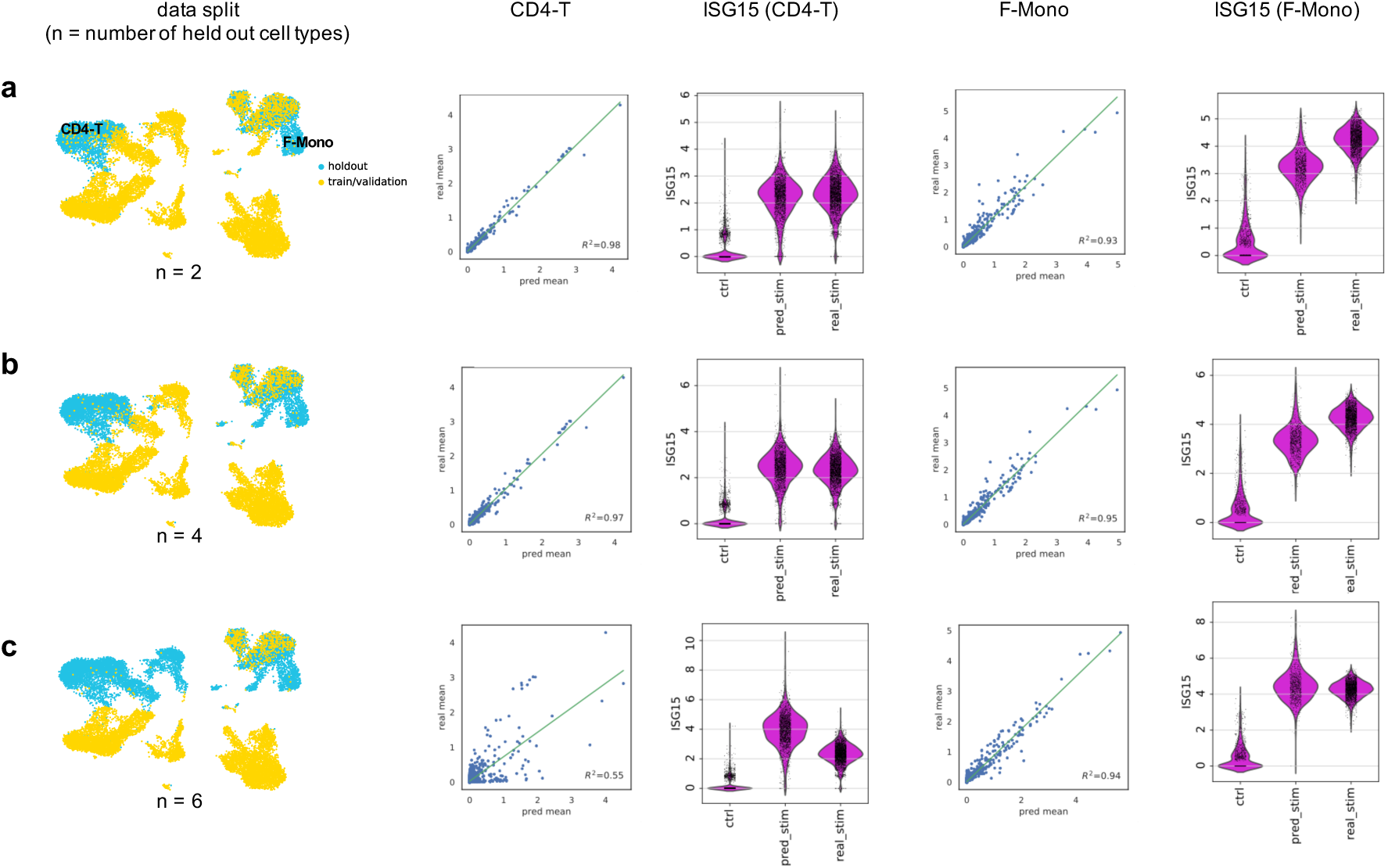
scGen performs robustly when holding out more than one cell type. **a-c,** Predicting IFN-*β* stimulated CD4-T and F-Mono cells form Kang *et al.* dataset in different scenarios with different number of held out cell types. First panel shows UMAP visualization for the position of held out cells. Other panels show mean gene expression of all genes and violin plot for ISG15, the top response gene after stimulation with IFN-*β* for CD4-T and F-mono cells.

#### Supplemental Note 7: Training and technical details

We used a similar architecture to train all models in all scenarios. This architecture includes reducing input dimension to 800 and creating another 800 features from the previous layer and finally projecting into 100 dimensional Gaussian governed latent space (*input*_*dim*_ ⟶ 800 ⟶ 800 ⟶ 100). The batch normalization [57] was applied to every layer except Gaussian and output layers. Leaky ReLU (Rectified Linear Unit) activation function was used for all the layers except Gaussian and output layers which linear and ReLU were used, Respectively. In order to avoid over-fitting, we exploited several techniques including dropout [58], *L*_2_ regularization and early-stopping. Note that, the degree of regularization, dropout rate, and early stopping hyper-parameters are the only changes we made to train the model on different datasets. Adam [59] optimizer with learning rate 0.001 was used to train the networks. The detailed hyper-parameters for each dataset are listed on the GitHub repository.

Usually, the conditions sizes are not equal leading to a biased *δ* vector estimation. Moreover, White [55] discovered that by removing smile vector from woman face, the male attribute was also added. This originates from the sampling bias induced by unequal size of smiling man and woman samples. In order to prevent a similar problem, as previously described we balanced cell type and condition size before estimating *δ*. Supplemental Figure 11 depicts the effects of using biased and unbiased *δ* vector for the prediction of stimulated CD4-T from Kang *et al.*

**Supplemental Figure 7.**
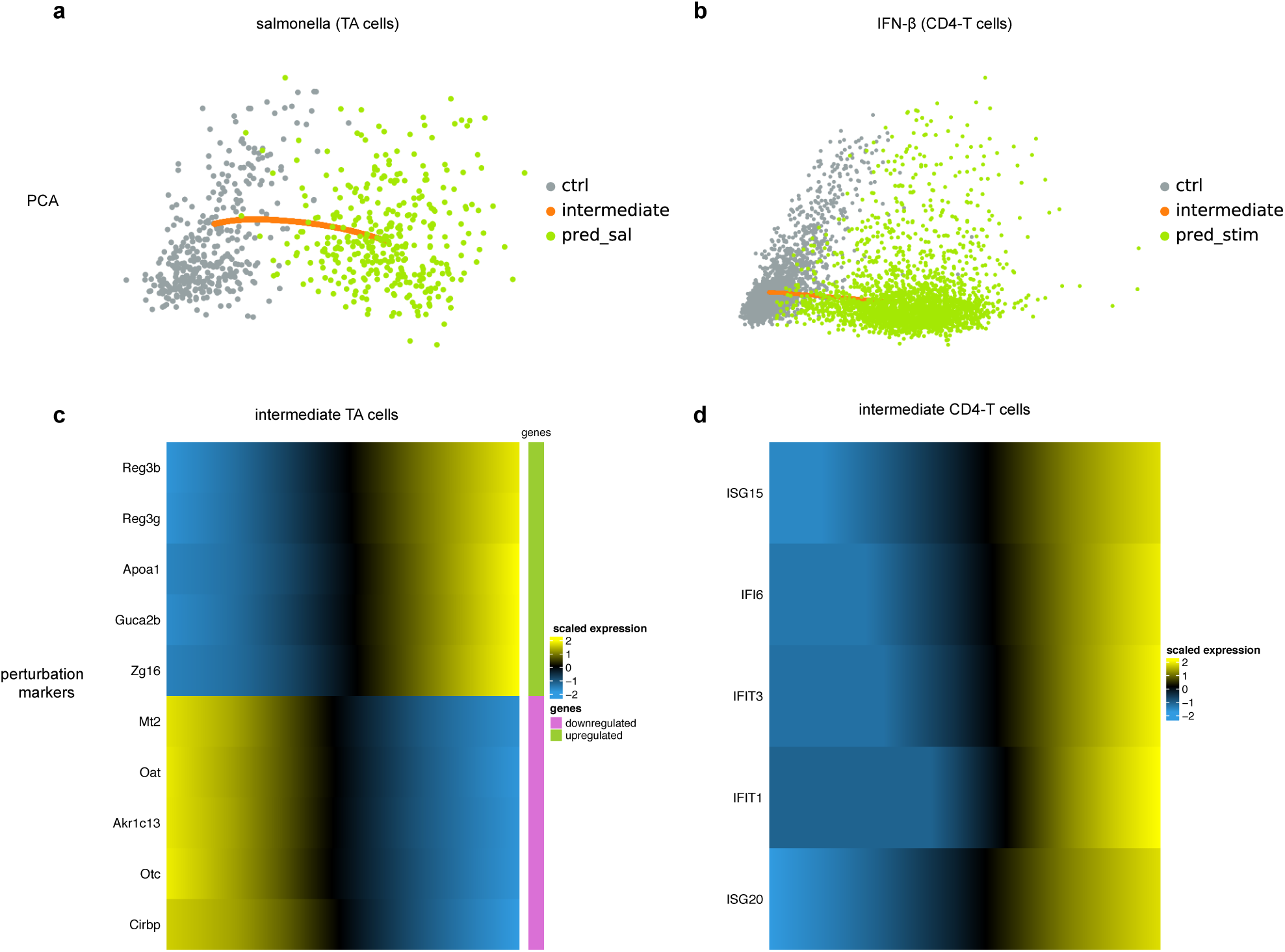
scGen enables the generation of intermediary cells between two conditions. **a-b,** PCA visualization of generated intermediary TA (Haber *et al.*) and CD4-T (Kang *et al.*) cells between control and predicted cells. **c,** Top five up and downregulated genes as we move from control to *Salmonella* infected cells. **d,** Similarly, variation of top five IFN-*β* marker genes while transitioning from control to predicted IFN-*β* stimulated cells.

#### Supplemental Note 8: Evaluations

**Silhouette width,** we calculated the Silhouette width based on the first 50 PCs of the corrected data or the latent space of the algorithm if it did not return corrected data. The Silhouette coefficient for for cell *i* is defined as:

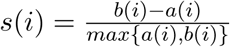

where *a*(*i*) and *b*(*i*) indicate the mean intra-cluster distance and the mean nearest-cluster distance for sample i, respectively. Instead of cluster labels one can use batch labels to asses batch correction methods. We used *silhouette_score* function from scikit-learn [60] to calculate the average Silhouette width over all samples.

**Error bars,** were computed by re-sampling the data points with replacement for 100 times and fitting the regression line for the re-sampled data. The interval represents the original estimation of *R*^2^ plus/minus the standard deviation of *R*^2^ values obtained from 100 fitted lines.

**cosine similarity,** computes the similarity as the normalized dot product of X and Y defined as:

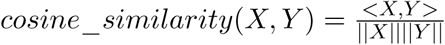

The *cosine_similarity* function from scikit-learn was used to compute cosine similarity.

**Supplemental Figure 8.**
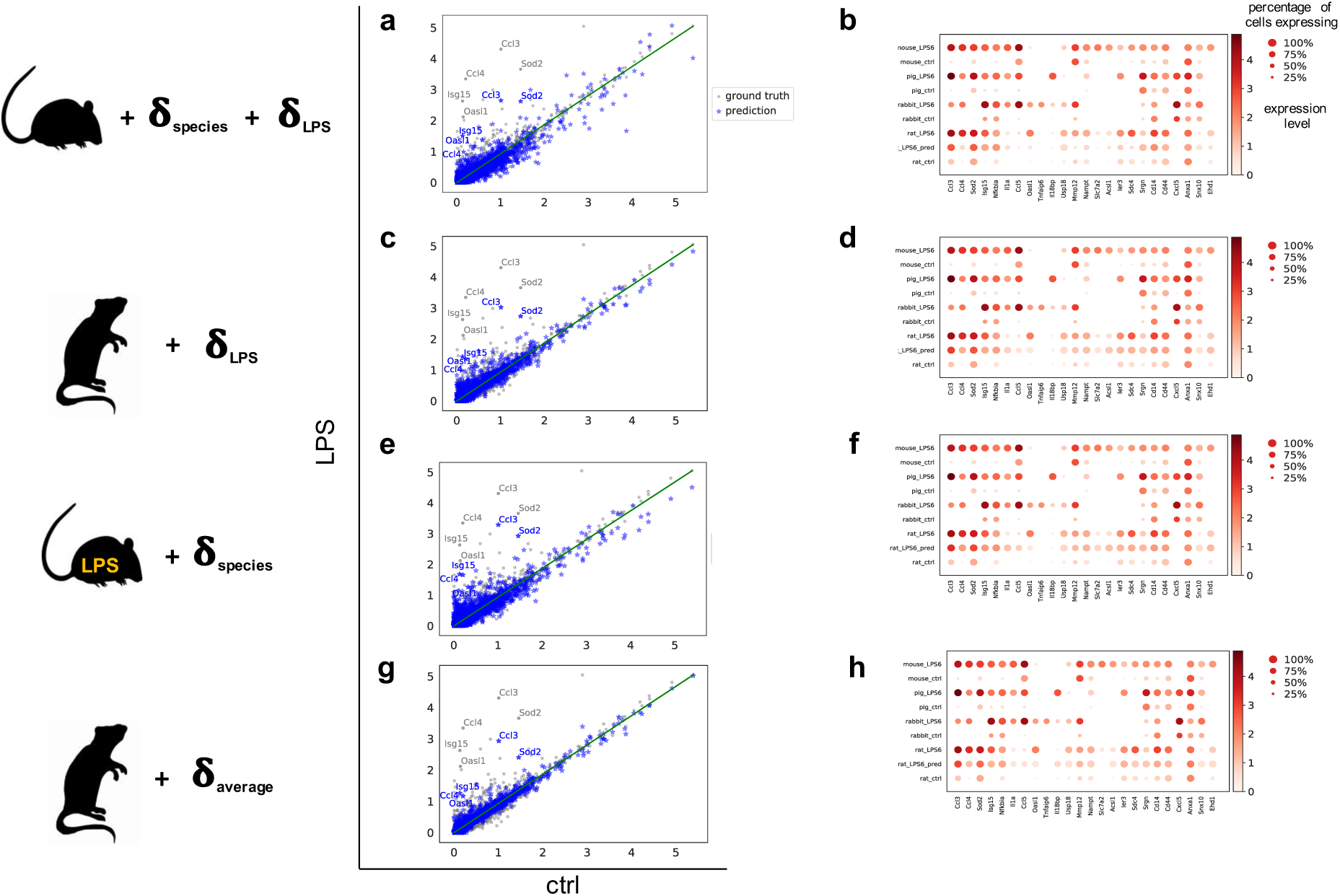
Alternative vector arithmetics for cross species prediction. **a-f,** Prediction of rat_LPS_ by adding difference vectors estimated using rat and mouse. **g-h,** Prediction of rat_LPS_ by adding *δ*_average_ to rat_control_ where *δ*_average_ = avg(*z*_LPS, all species_) − avg(*z*_control, all species_).

**Supplemental Figure 9.**
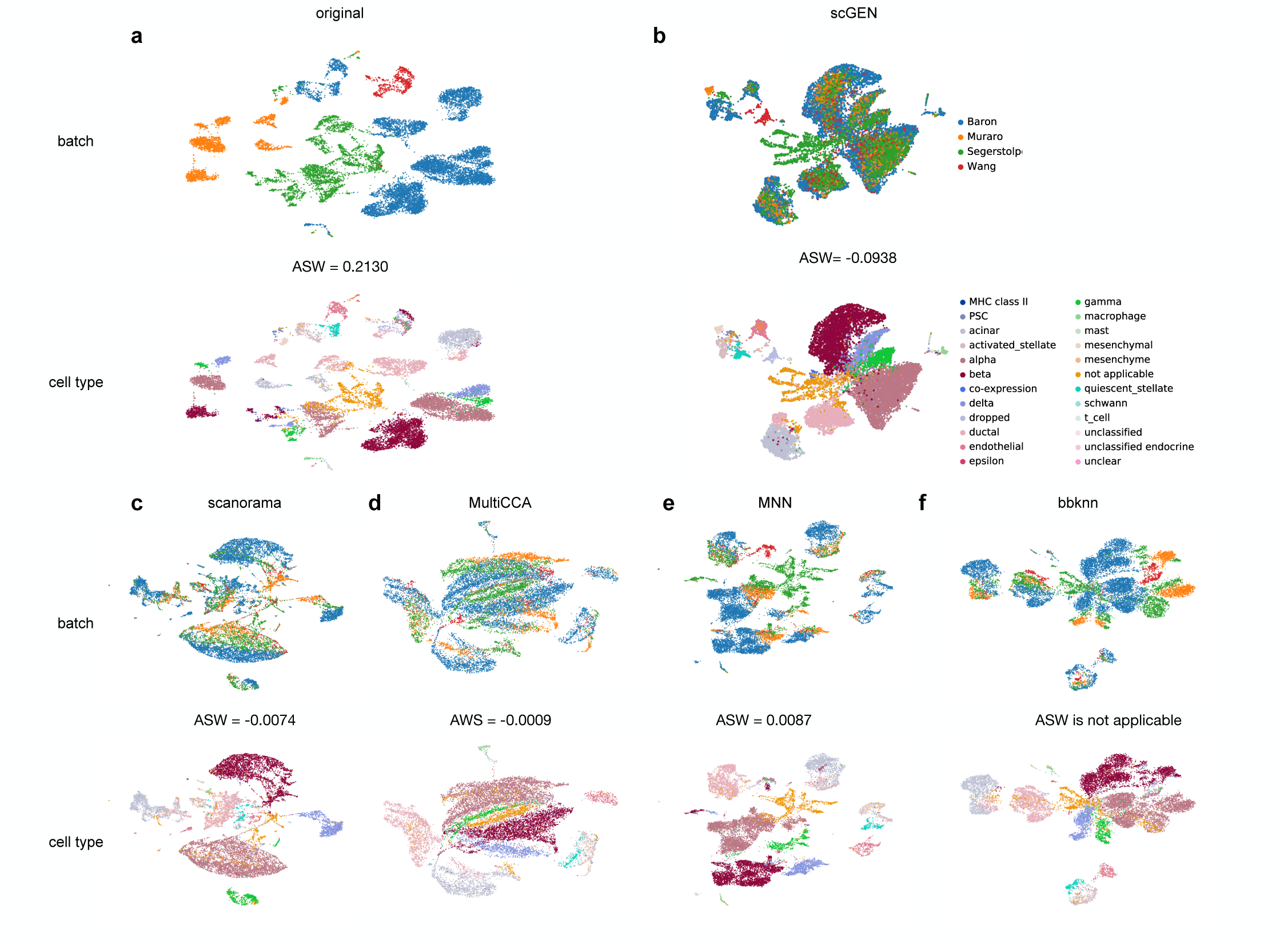
Comparison of existing batch effect removal methods at integrating four different pancreatic datasets. **a,** Original data contains large technical variation which causes similar cell types cluster separately. We report average silhouette width (ASW) for batches in the original data (ASW = 0.2130, lower is better). **b,** scGen aligns shared cell types in different studies while preserving study specific cell types independent after batch correction and returns lowest ASW (−0.0938). **c,** Scanorama marges shared cell types but they are not perfectly mixed and does not persevere the structure of the small study specific cell types. **d,** CCA connects batches well but shared cell types are not perfectly mixed. **e,** MNN mixes some cell types while keeping batch effect for others and it successfully preserves structure of study specific cell types. **f,** Results of bbknn show shared cell types are not perfectly mixed and some cell types are mistakenly merged into wrong clusters. In contrast to other methods this model only returns modified KNN graph and does not provide any form of corrected data thus ASW is not directly applicable to corrected data.

**Supplemental Figure 10.**
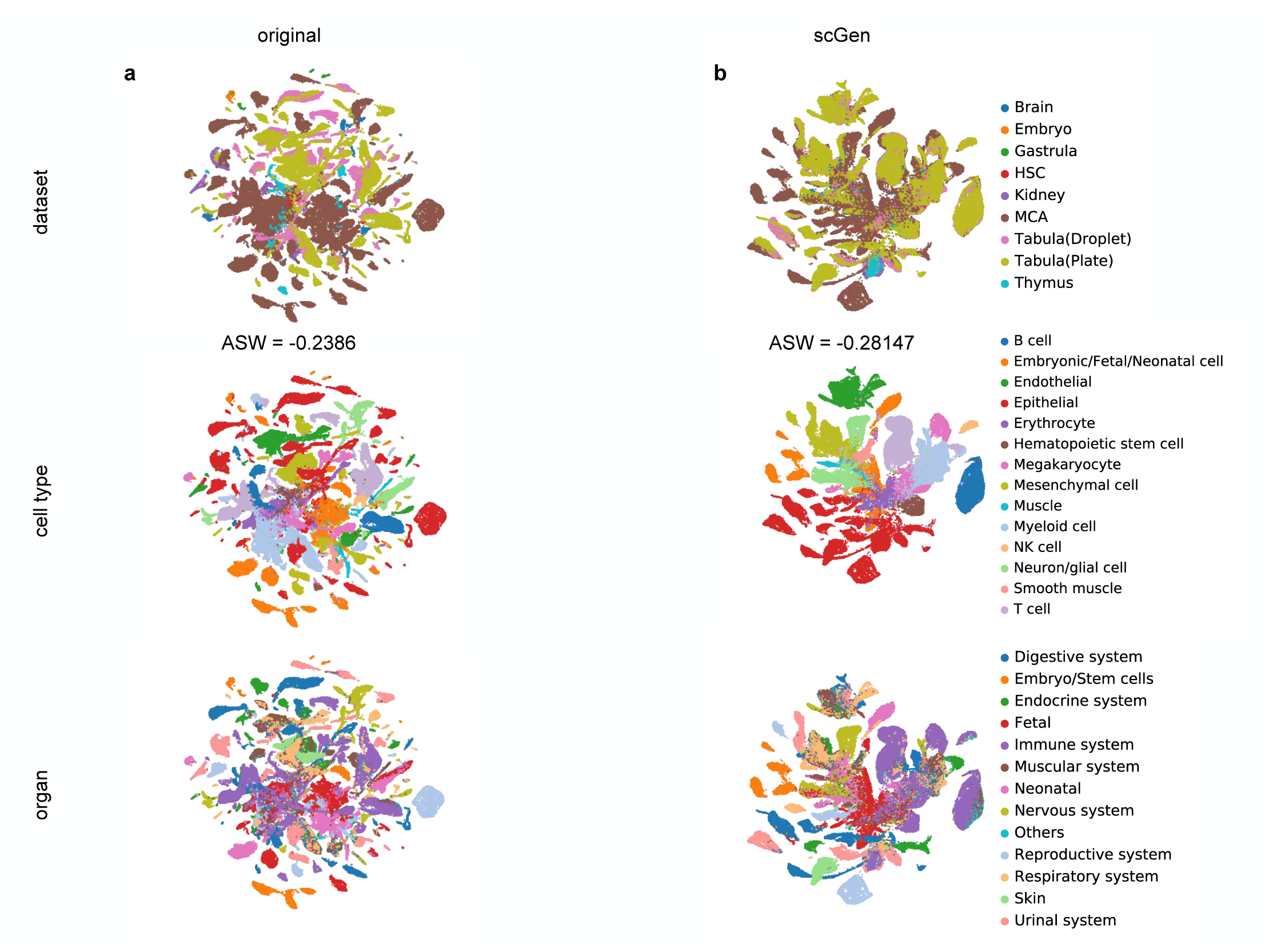
scGen integrates eight mouse single cell atlasess with 114600 cells. **a,** UMAP visualization of eight different datasets with their corresponding study, cell type and organ labels. ASW was calculated based on the 57300 randomly sub-sampled cells with their study labels. **b,** scGen merges the data by connecting the similar cell types according to their cell labels while having lower ASW (−0.28147).

**Supplemental Figure 11.**
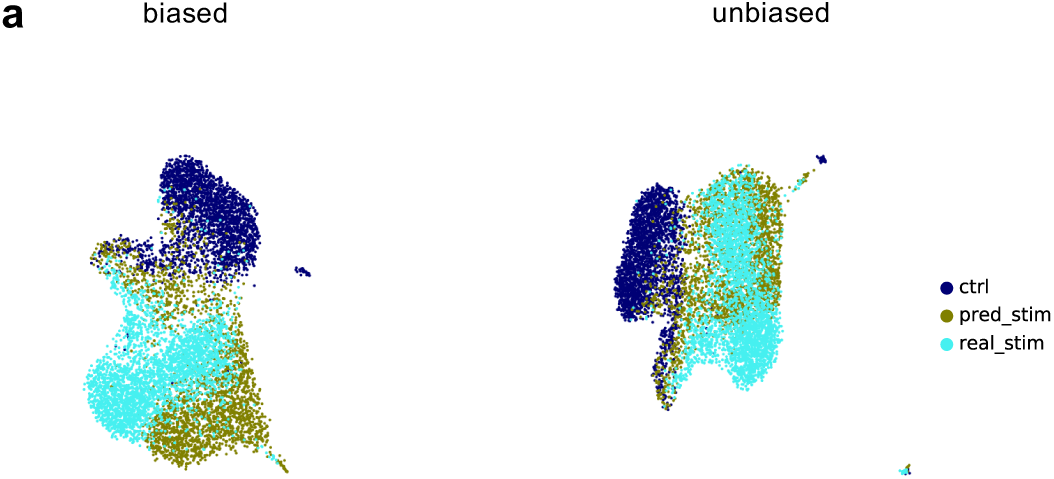
Biased sampling effect. **a,** UMAP visualization of CD4-T cells prediction depicts that unbiased predicted cells have more overlap with real stimulated cells than biased predictions.

